# Establishment of a stable *zg6pd*^*M118-144*^ transgenic zebrafish model of glucose-6-phosphate dehydrogenase deficiency

**DOI:** 10.1101/2020.04.28.066779

**Authors:** Hai-Xiong Xia, Yan-Hua Zhou, Yuan-Yuan Tuo, Ping-Ping Ren, Jin Song, Lu-Jun Shang, Jiao Jin, Zhi-Xu He, Li-Ping Shu

**Affiliations:** National Guizhou Joint Engineering Laboratory for Cell Engineering and Biomedicine Technique, State Key Laboratory of Functions and Applications of Medicinal Plants, Guizhou Province Key Laboratory for Regenerative Medicine, Department of Immunology, Department of Pediatrics, Guizhou Medical University, 9 Beijing Road, Guiyang 550004, People’s Republic of China; Key Laboratory of Adult Stem Cell Transformation Research, Chinese Academy of Medical Sciences, China; Department of Pediatrics, Zunyi Medical University, 201 Dalian Road, Zunyi 560004, China; Jinyang hospital of Guiyang, 547 Jinyang South Road, Guiyang 550023, China

**Author notes:** These three authors contributed equally to this work (Hai-Xiong Xia, Yan-Hua Zhou, Yuan-Yuan Tuo). Correspondence to: Dr. Li-Ping Shu, PhD. Tissue Engineering and Stem Cell Research Center, Guizhou Medical University, 9 Beijing Road, Guiyang 550004, People’s Republic of China.; And Dr. Zhi-Xu He, PhD. Tissue Engineering and Stem Cell Research Center, Guiyang Medical University, 9 Beijing Road, Guiyang 550004, People’s Republic of China.

## Abstract

Glucose-6-phosphate dehydrogenase (G6PD) deficiency, the most common genetic defect and enzymopathy with a wide distribution and increased public health concern, predisposes subjects succumb to oxidative stress. G6PD deficiency has been associated with hemolysis. Clinically, G6PD deficiency is asymptomatic and the clinical manifestations occur with the exposure to certain agents. Due to the lack of suitable animal models that can predict the clinical hemolytic potential of drugs, it needs an appropriate research model to fully recapitulate the manifestations of G6PD deficiency in clinic, to optimize the malaria therapy and promote anti-malarias development. The present study has displayed a stable transgenic *Tg*(*zgata1-g6pd*^*M118-144*^-*egfp*) zebrafish model with G6PD deficiency which mimics the clinical features of G6PD deficiency phenotypically and functionally. The findings showed that there was an inadequate level of reduced GSH in the transgenic *Tg*(*zgata1-g6pd*^*M118-144*^-*egfp*) zebrafish line in the presence or absence of α-naphthol, compared to the wildtype zebrafish, indicating an attenuation of g6pd activity in the transgenic zebrafish line. In addition, the observations show that there is a less abundance of g6pd in the transgenic *Tg*(*zgata1-g6pd*^*M118-144*^-*egfp*) zebrafish line. On the other hand, there is no morphological abnormality in the transgenic *Tg*(*zgata1-g6pd*^*M118-144*^-*egfp*) zebrafish line. Taken together, our work has delivered a novel stable transgenic zebrafish model of G6PD deficiency that will facilitate the mechanistic and functional elucidation for the role of G6PD in erythrocytic pathophysiology. This model will promote the translational research for the drug development, in particular, for anti-malarias development.

## Introduction

Glucose-6-phosphate dehydrogenase (G6PD), a cytosolic enzyme (EC:1.1.1.49), possesses a vital role in the regulation of redox homeostasis and erythrocytic pathophysiology [1]. Human G6PD is encoded by a housekeeping X-linked gene at the Xq28 locus, spanning 18 kb with 13 exons; and human G6PD consists of 515 amino acids [2]. G6PD mainly functions to generate nicotinamide adenine dinucleotide phosphate (NADPH) [3, 4]. It is the first and the rate-limiting enzyme in the pentose phosphate pathway (PPP), oxidizing glucose-6-phosphate to 6-phosphogluconolactone and reducing NADP^+^ to NADPH. NADPH serves as a key electron donor and keeps glutathione in its reduced form in the defense against oxidizing agents and in reductive biosynthetic reactions, so as to detoxify dangerous oxidative metabolites maintain redox homeostasis [3]. Therefore, attenuated G6PD activity succumbs cells to oxidative stress [3].

G6PD deficiency, the most common X-linked enzymopathy in humans, affects about 400 million people worldwide in malaria endemic areas, with male being primarily affected [5–7]. It is particularly high prevalent in tropical Africa, the Middle East, tropical and subtropical Asia, some areas of the Mediterranean, and Papua New Guinea [5, 6]. The prevalence of G6PD deficiency is ~10% in US, with black males being mainly affected [5]. G6PD deficiency mainly affects Chinese subjects in the South areas [8]. Conventionally, inherited G6PD deficiency is asymptomatic [7]. The clinical manifestations of G6PD deficient subjects include acute episodic hemolytic anemia, chronic hemolytic anemia, and kernicterus when exposed to sulfonamide antibiotics, antimalarials, and fava beans [7, 9]. These pro-oxidative or oxidative drugs or chemicals inflict an oxidative stress on cells, resulting in a disruption in cellular biological events and cellular damages [3, 10].

Hemolysis is the most common consequence for G6PD deficient subjects when exposed these agents due to oxidative stress [9, 11]. G6PD controls PPP that is the only source for NADPH in RBCs, justifying the singularly important role of G6PD in NADPH generation for the maintenance of redox homeostasis in RBCs [1, 3]. Thus, RBCs with attenuated G6PD activity are more vulnerable to oxidative stress and G6PD deficiency predispose subjects to oxidative stress induced damages, such as RBC destruction and hemolysis, when exposed to oxidative agents. However, due to the asymptomatic feature of G6PD deficiency, it hinders the utilization of drugs for malaria treatment in clinical practice. Also, the lack of appropriate animal models of G6PD deficiency stymies the development of efficacious anti-malaria agents. Thus, there is a great demand for suitable G6PD deficient model so as to facilitate and promote the mechanistic and translational research on G6PD deficiency. Previously, mouse models G6PD deficiency have been developed to mimicking G6PD deficiency in humans, but it is limited due to the early embryonic death and lack of significant response to oxidative challenge [12]. On the other hand, zebrafish has increased popularity as a model for human diseases [13], thanks to the merit of external development and optical clarity to visualize cellular events *in vivo*. Recently, a zebrafish model of G6PD deficiency was established using morpholinos targeting the 5-prime exons of *g6pd* [14]. Although it is limited by the transient knockdown of *g6pd*, it provides a clue to develop zebrafish model with G6PD deficiency.

Due to the lack of suitable animal models that can predict the clinical hemolytic potential of drugs, the present study aimed to establish a stable transgenic zebrafish model with G6PD deficiency, to examine the role of G6PD in erythrocytic pathophysiology and its potential as a therapeutic target in the regulation of redox homeostasis, cell growth, and embryonic development.

## Methods

### Zebrafish strain and fish husbandry

All zebrafish lines were housed in Guiyang Medical University Zebrafish Core Facility and the breeding and staging of zebrafish lines were performed as described in our previous study [13]. The protocol was approved by the Animal Experimentation Committee of Guiyang Medical University, Guizhou, People’s republic of China. *Tg:zgata1:egfp* zebrafish line was a precious gift from Dr. Min Deng (Institute for Nutritional Sciences, SIBS, Chinese Academy of Sciences, Shanghai, China).

### Primer design

To generate the mutant zebrafish g6pd with the deletion of amino acid residues from 40 to 48, the nucleotide was deleted from 118 to 144. Two pairs of primer for *zg6pd*^*M118-144*^ were designed according to zebrafish cDNA (NCBI Reference Sequence: XM_694076.7). The sequences are listed in Table 1.

**Table 1.**
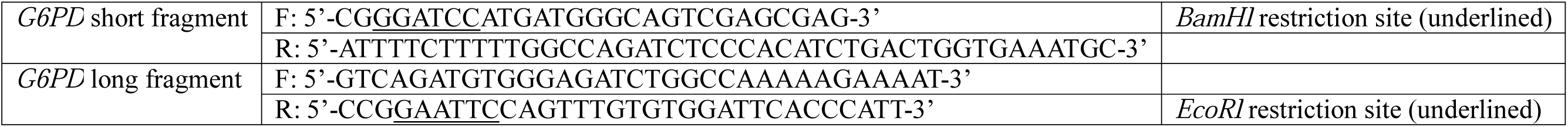
The sequences for the *G6PD* primers.

### Constructs

The recombinant plasmids were constructed as previously described [13]. In brief, the plasmids were constructed via *BamHI* and *EcoRI*-mediated digestion and T4 DNA ligase-mediated ligation according to the manufacturer’s instruction (Fermentas, Canada) (Table 2). All recombinant plasmids were sequenced to ensure that the plasmids were correctly constructed.

**Table 2.**
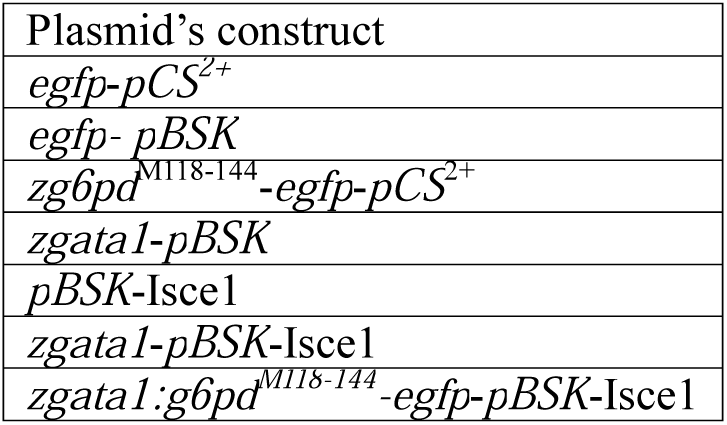
List of the plasmids used in this study.

### Micro-injection and generation of stable transgenic zebrafish lines

The micro-injection was performed to generate stable transgenic zebrafish lines as previously described [13, 15]. The potential founders (F0) were bred with wildtype fish and embryos were screened for green fluorescent protein (GFP) expression at 20–24 hours after fertilization (hpf). Those showing tissue-specific GFP expression were grown to adulthood (F1). The homozygous transgenic zebrafish line (F2) was descended from F1. Due to the possibility of mosaic integration in the germline, potential founders were only scored as non-transgenic after at least 100 of their offspring were shown to be negative for GFP expression.

### Microscopic imaging

To examine the expression of *g6pd*^*M118-144*^, the micro-injected embryos with *zgata1:g6pd*^*M118-144*^-*egfp*-*pBSK-I-SceI* plasmids were imaged as previously described [13]. The embryos were screened for GFP expression using a Zeiss SteREO Discovery V12 fluorescence stereomicroscope. The images were acquired with the same microscope equipped with an AxioCam MRC5 digital camera and analyzed by AxioVision software.

### Western blotting analysis

The proteins samples were collected as previously described [13]. Briefly, the embryos were deyolked and homogenized. The harvested lyses were prepared for fragmentation on SDS-polyacrylamide gel. The corresponding antibodies were used to evaluate the expression level of EGFP and G6PD.

### Genotyping and DNA sequencing

Genomic DNA was collected and used for genotyping. The PCR conditions were as follows: 94ºC for 8 min; 35 cycles of 94ºC for 40 sec; 65ºC for 30 sec; 68ºC for 90 sec; and then 68ºC for 15 min. The resultant products were collected for further DNA sequence (Sunnybio, Shanghai, China). Genomic DNA from embryos of wildtype and *zgata1-egfp* zebrafish line were used as control.

### *In situ* hybridization

Digoxin- and fluorescein-labeled *egfp* RNA probes were used for the whole-mount mRNA *in situ* hybridization (WISH) assay to examine the expression of *g6pd*^*M118-144*^ as previously described [13]. Embryos were imaged with an AxioCam MRC5 digital camera and analyzed by AxioVision software. Embryos of wildtype and *zgata1-egfp* zebrafish line were used as control.

### G6pd enzyme activity

G6pd activity was determined as previously described with some modifications [16, 17]. Briefly, g6pd activity was evaluated in embryos in the presence or absence of α-naphthol (10 µM) over 48 h via the measurement of G6PD/6PGD ratio. The absorbance was measured at 650 nm. Also, g6pd activity was evaluated using fluorescence spot test as previously described [18, 19]. In brief, the embryos were exposed to α-naphthol (10 µM) over 48 h and the activity was measured via the visual evaluation of fluorescence reduced NADPH when activated by UV light at 360 nm. The fluorescence of NADPH is proportional to the activity of g6pd.

### GSH measurement

The GSH level was measured using reduced glutathione assay kit according to the manufacturer’s instruction (Nanjing Jiancheng Bioengineering Institute, Nanjing, China). In brief, the wildtype and transgenic embryos at 24 hpf were exposed to α-naphthol at 5, 10, and 20 µM for 48 h and harvested at 72hpf. The supernatant was collected for GSH detection in 96-well plates and the absorbance was measured at 405 nm.

### Cardiovascular toxicity evaluation

The cardiovascular toxicity was evaluated through the assessment of pericardium malformation rate in the F3-generation homozygous transgenic zebrafish line. The F3-generation embryos were exposed to α-naphthol at 5, 10, and 20 µM for 48 h and primaquine at 150 and 200 µM for 72 h. The zebrafish lines were visualized and the images were acquired at indicated phenotypic endpoints under the dissecting stereomicroscope.

### Hemoglobin staining & Erythrocyte staining

Hemoglobin staining was performed as described previously [20]. The dechorionated live embryos were stained in 0.6 mg/mL o-dianisidine staining system and dehydrated through graded ethanol washes of 25%, 50%, and 100%. The processed embryos were subject to visual observation using Nikon stereomicroscope with NES imaging system. The images were analyzed using Image J. On the other hand, erythrocyte staining was performed as previously described [21]. The erythrocytes were collected and stained with Wright-Giemsa or Prussian blue [22]. Images were obtained using NES system under Zess microscope.

### Flow cytometry

The apoptosis of erythrocyte was determined using PE annexin-V apoptosis detecting kit according to the manufacturer’s instruction (BD, USA). In brief, the embryos were exposed to α-naphthol at 5, 10, and 20 µM for 48 h and harvested at 72hpf. The erythrocytes were collected and stained with annexin-V and 7-AAD solution for 15 min in the dark. Subsequently, the cells samples were subject to flow cytometry (Beckman FC500)

### Statistical analysis

Data are expressed as mean ± SD. One-way analysis of variance (ANOVA) followed by Tukeys multiple comparison procedure was used for comparisons of multiple groups. P<0.05 was considered to be statistically significant. The assays were performed at least three times independently.

## Results

### Construction of *zgata1-g6pd*^*M118-144*^-*egfp-pBSK-I-SceI* plasmid

We constructed a *zgata1-g6pd*^*M118-144*^-*egfp-pBSK-I-SceI* plasmid with the nucleotide deletion from 118 to 144 through *BamHI* and *EcoRI*-mediated digestion and T4 DNA ligase-mediated ligation (Figure 1A). There were four fragmentations that were visualized with an approximate size of 5 533, 3 227, 1 539, and 717bp after digested by *BamHI*, *EcoRI*, and *XhoI* (Figure 1B). These four fragmentations respectively indicated *gata1* promoter, *pBSK-I-SceI* block, *g6pd*^*M118-144*^, and *egfp*. This result clearly showed the correct construction of *zgata1-g6pd*^*M118-144*^-*egfp-pBSK-I-SceI* plasmid. Furthermore, the fragmentations were sequenced and the sequence provided further proof for the construction of *zgata1-g6pd*^*M118-144*^-*egfp-pBSK-I-SceI* plasmid (Figure 1C).

### Expression profile of *zgata1-g6pd*^*M118-144*^-*egfp-pBSK-I-SceI* plasmid in zebrafish

We next performed micro-injection of *zgata1-g6pd*^*M118-144*^-*egfp-pBSK-I-SceI* plasmid into zebrafish embryos to examine the expression pattern of *g6pd*^*M118-144*^ in wildtype zebrafish via the visualization of egfp *in vivo*. Egfp was observed at 11hpf at anterior lateral mesoderm and posterior lateral mesoderm and in blood circulation system at 24hpf. The green fluorescence signal decayed as age. No green fluorescence was detected in wildtype zebrafish without micro-injection (Figure 2). A total number of 9 800 embryos that were micro-injected with *zgata1-g6pd*^*M118-144*^-*egfp-pBSK-I-SceI* plasmid. The green fluorescence was observed in 42 zebrafish, of which 5 F0 generation zebrafish were selected with stable inheritance of egfp in F1 generation as shown in Figure 3A and B. The expression pattern of g6pd was monitored over 72 h, which overlapped the expression pattern of *gata1*. The expression pattern of g6pd was similar to that in control zebrafish with micro-injection of *Tg(zgata1-egfp)* plasmid (Figure 3B).

### Verification of *g6pd*^*M118-144*^ expression in F1

Subsequently, we verified the expression of *g6pd*^*M118-144*^ in F1 generation transgenic zebrafish. The genomic DNA was extracted from F1 at 5-day post fertilization and was subject to genomic PCR for the examination of*g6pd*^*M118-144*^ expression. The resultant products were fragmented and a clear band with an approximate size of 1 539bp was visualized, indicating the fragmentation of *g6pd*^*M118-144*^ (Figure 4A). Also, the resulting fragmentation was recovered and sequenced. The alignment indicated that the sequence of the fragment was identical to that in *zgata1-g6pd*^*M118-143*^-*egfp-pBSK-I-SceI* plasmid (Figure 4B).

### Identification the protein expression in F1

To further corroborate the expression of g6pd in the stable transgenic zebrafish line, WISH assay was employed to examine the expression of *egfp*. The expression of *egfp* was detected in wildtype, *Tg*(*zgata1-egfp*), and *Tg*(*zgata1-g6pd*^*M118-144*^-*egfp*) zebrafish lines using digoxin labeled *egfp* anti-sence mRNA probe. The *in situ* hybridization results showed that *egfp* was expressed at intermediate cell mass in both *Tg*(*zgata1-egfp*) and *Tg*(*zgata1-g6pd*^*M118-144*^-*egfp*) zebrafish lines and at dorsal vessel only in *Tg*(*zgata1-egfp*) zebrafish line. No detectable *egfp* mRNA level was observed in wildtype zebrafish line (Figure 5A). The protein expression level of egfp was examined. As shown in Figure 5B, egfp was abundantly expressed in *Tg*(*zgata1-egfp*) and *Tg*(*zgata1-g6pd*^*M118-144*^-*egfp*) zebrafish lines, evident from the band at 27 kDa. Of importance, g6pd-egfp fusion protein was detected in *Tg*(*zgata1-g6pd*^*M118-144*^-*egfp*) zebrafish line as indicated by the band at 88 kDa probed with egfp primary antibody. Taken together, the *zgata1-g6pd*^*M118-143*^-*egfp-pBSK-I-SceI* plasmid was able to expressed and stably inherited in zebrafish.

### G6PD activity in transgenic zebrafish line

The subsequent functional assays were performed to evaluate the phenotypic alterations in g6pd deficient zebrafish. First, the g6pd activity was examined in F1 heterozygous and F3 homozygous *Tg*(*zgata1-g6pd*^*M118-144*^-*egfp*) transgenic zebrafish line. As shown Figure 6A, no significant alteration in g6pd activity was observed in F1 heterozygous *Tg*(*zgata1-g6pd*^*M118-144*^-*egfp*) transgenic zebrafish in the presence or absence of 10 µM α-naphthol, compared to the wildtype zebrafish. Also, no dramatic change was observed in g6pd activity in embryos for F3 homozygous *Tg*(*zgata1-g6pd*^*M118-144*^-*egfp*) transgenic zebrafish line (g6pd^M118-144^-3) (Figure 6B). These asymptomatic features reflect the clinical manifestations of G6PD deficiency. Given the vital role of g6pd in the maintenance of the reducing power in cells, in particular, the RBCs [7], we following examined the reduced GSH level in F3 homozygous *Tg*(*zgata1-g6pd*^*M118-144*^-*egfp*) transgenic zebrafish line (g6pd^M118-144^-3). As would be expected, in comparison to the wildtype zebrafish, there was a significant decrease in the level of reduced GSH in F3 transgenic zebrafish with or without exposure to α-naphthol (Figure 6C and D). Meanwhile, there was no abnormality observed in the F3 homozygous *Tg*(*zgata1-g6pd*^*M118-144*^-*egfp*) transgenic zebrafish line (g6pd^M118-144^-3) (Figure 6E). In aggregate, these results show that g6pd deficient zebrafish line further recapitulates the clinical features in G6PD deficient subjects.

### Cardiovascular toxicity in F3 homozygous *Tg*(*zgata1-g6pd*^*M118-144*^-*egfp*) transgenic zebrafish exposed to primaquine and α-naphthol

At last, we performed assays to evaluate cardiovascular toxicity-associated morphological abnormalities of F3 homozygous *Tg*(*zgata1-g6pd*^*M118-144*^-*egfp*) transgenic zebrafish line. As shown in Figure 7A and B, there was no morphological changes that were found in the heart malformation in F3 homozygous *Tg*(*zgata1-g6pd*^*M118-144*^-*egfp*) transgenic zebrafish line (g6pd^M118-144^-3) in the presence or absence of primaquine or α-naphthol. The hemoglobin and erythrocyte stainings also showed no significant alteration in hemolysis and no abnormality was observed in erythrocyte (Figure 7C and D). Furthermore, the specific cells expressing egfp were collected and subject to apoptosis assay. The results did not show significant difference in apoptosis of erythrocytes in g6pd^M118-144^-3 zebrafish line in the presence or absence of α-naphthol, compared to the *Tg*(*zgata1-egfp*) transgenic zebrafish (Figure 7E).

## Discussion

G6PD deficiency is the most common genetic defect and enzymopathy with a wide distribution and increased public health concern. G6PD deficiency subjects succumb to oxidative stress due to lack of reducing power in response to oxidative challenges. G6PD deficiency has been associated with hemolysis. Clinically, G6PD deficiency is asymptomatic and the clinical manifestations occur with the exposure to certain agents, such as anti-malarias. In order to optimize the malaria therapy and promote anti-malarias development, it therefore needs an appropriate research model to fully recapitulate the manifestations of G6PD deficiency in clinic. The present study has displayed a stable transgenic zebrafish model with G6PD deficiency which mimics the clinical features of G6PD deficiency phenotypically and functionally.

G6PD deficiency is polymorphic and over 400 *G6PD* polymorphisms have been identified. There are a number of functionally important polymorphisms that have been identified, such as A95G, G392T, G487A, A493G, C592T, C1024T, C1360T, G1376T, and G1388A [23]. These nucleotide substitutions lead to alteration in structure and activity [24, 25]. The consequence of nucleotide deletion from 118 to 144 would be expected that the deletion would structurally affect the dimerization of G6PD, which in turn compromises the function and activity of G6PD. Of note, the transgenic *Tg*(*zgata1-g6pd*^*M118-144*^-*egfp*) zebrafish line did not show phenotypic abnormality, compared to the wildtype. It suggests that this transgenic model phenotypically mimics the clinical features of G6PD deficiency. This construction would provide a novel G6PD deficient model for future mechanistic and functional studies.

Functionally, G6PD is the first and the rate-limiting enzyme in PPP for NADPH generation [3]. NAPDH is critical for many essential cellular systems and fueling GSH pool in response to oxidative challenges [1]. Because erythrocytes do not produce NADPH in any other way than PPP, they are more easily succumb to oxidative stress and more susceptible than any other cells to oxidative damages. It renders the singularly important role of G6PD in erythrocytic pathophysiology. Recently, a number of studies have revealed the important role G6PD in erythrocytic pathophysiology, such as redox homeostasis and embryonic development [1]. Also, G6PD has been considered as a promising therapeutic targets for enzymopathy and beyond [11, 26, 27]. In the present study, the findings show that there is an inadequate level of reduced GSH in the transgenic *Tg*(*zgata1-g6pd*^*M118-144*^-*egfp*) zebrafish line in the presence or absence of α-naphthol, compared to the wildtype zebrafish, indicating an attenuation of g6pd activity in the transgenic zebrafish line. In addition, the observations show that there is a less abundance of g6pd in the transgenic *Tg*(*zgata1-g6pd*^*M118-144*^-*egfp*) zebrafish line, which may be ascribed to the deletion-induced protein instability and degradation. It, in turn, results in the decline in the activity. On the other hand, there is no morphological abnormality in the transgenic *Tg*(*zgata1-g6pd*^*M118-144*^-*egfp*) zebrafish line.

Taken together, our work has delivered a novel stable transgenic zebrafish model of G6PD deficiency that will facilitate the mechanistic and functional elucidation for the role of G6PD in erythrocytic pathophysiology. This model will promote the translational research for the drug development, in particular, for anti-malarias development.

## Acknowledgement

This project was supported in part by the National Natural Science Foundation of China (31860325, 31360285), and in part by the Guizhou Province’s Science and Technology Major Project (Qian-P-Ren[2017]5611 & Qian-P-Ren[2019]5406), and in part by the Non-profit Central Research Institute Fund of Chinese Academy of Medical Sciences(NO.2018PT31048, 2019PT310013).

## Conflict of interest

The authors declare that there is no conflict of interest.

---

Figure 1. Construction of *zgata1-g6pd*^*M118-144*^-*egfp-pBSK-I-SceI* plasmid. A: Map of *zgata1-g6pd*^*M118-144*^-*egfp-pBSK-I-SceI* plasmid. B: Electrophoresis of *zgata1-g6pd*^*M118-144*^-*egfp-pBSK-I-SceI* plasmid fragmentations after BamHI, EcoRI, and XhoI mediated digestion. Lane 1: DNA marker, land 2: *zgata1-g6pd*^*M118-144*^-*egfp-pBSK-I-SceI* plasmid digestion products, lane 3: blank control. B: The sequence of *zgata1-g6pd*^*M118-144*^-*egfp-pBSK-I-SceI* plasmid.

Figure 2. Expression of *zgata1-g6pd*^*M118-144*^-*egfp-pBSK-I-SceI* plasmid in wildtype zebrafish. Expression profile of g6pd in wildtype zebrafishe with or *zgata1-g6pd*^*M118-144*^-*egfp-pBSK-I-SceI* plasmid at 11, 24, 30, and 36hpf.

Figure 3. Expression of *Tg*(*zgata1-g6pd*^*M118-144*^-*egfp*) in F1 generation of transgenic zebrafish. A: The flowchart of trangenic zebrafish breeding. B: The expression pattern of green fluorescence in F1 *Tg*(*zgata1-g6pd*^*M118-144*^-*egfp*) and *Tg*(*zgata1-egfp*) control trangenic zebrafish lines.

Figure 4. Expression of *Tg*(*zgata1-g6pd*^*M118-144*^-*egfp*) in F1 generation of transgenic zebrafish. A: The electrophoresis of *g6pd*^*M118-144*^ in *Tg*(*zgata1-g6pd*^*M118-144*^-*egfp*) F1 generation of transgenic zebrafish. B: The sequence of *g6pd*^*M118-144*^.

Figure 5. Identification the expression of egfp and g6pd in F1 generation transgenic zebrafish. A: Expression of *egfp* mRNA in F1 generation wildtype, *Tg*(*zgata1-egfp*), and *Tg*(*zgata1-g6pd*^*M118-144*^-*egfp*) zebrafish lines at 25hpf. B: Protein expression of egfp in F1 generation wildtype (lane 1), *Tg*(*zgata1-egfp*) (lane 2), and *Tg*(*zgata1-g6pd*^*M118-144*^-*egfp*) (lane 3) zebrafish lines.

Figure 6. G6pd activity and reduced GSH level in *Tg*(*zgata1-g6pd*^*M118-144*^-*egfp*) transgenic zebrafish. A: G6pd activity in F1 heterozygous *Tg*(*zgata1-g6pd*^*M118-144*^-*egfp*) transgenic zebrafish in the presence or absence of 10 µM α-naphthol. B:: G6pd activity in F3 homozygous *Tg*(*zgata1-g6pd*^*M118-144*^-*egfp*) transgenic zebrafish. C: Reduced GSH level in *Tg*(*zgata1-g6pd*^*M118-144*^-*egfp*) transgenic zebrafish. D: Reduced GSH level in *Tg*(*zgata1-g6pd*^*M118-144*^-*egfp*) transgenic zebrafish with exposure of α-naphthol at 5, 10, and 20 µM. E: Phenotype of F3 homozygous *Tg*(*zgata1-g6pd*^*M118-144*^-*egfp*) transgenic zebrafish (g6pd^M118-144^-3).

Figure 7. Cardiovascular toxicity-associated morphological abnormalities of F3 homozygous *Tg*(*zgata1-g6pd*^*M118-144*^-*egfp*) transgenic zebrafish line. Heart malformation rate in *Tg*(*zgata1-g6pd*^*M118-144*^-*egfp*) transgenic zebrafish in the presence of primaquine (A) and α-naphthol (B). Hemoglobin staining (C) and erythrocyte staining (D) for F3 homozygous *Tg*(*zgata1-g6pd*^*M118-144*^-*egfp*) transgenic zebrafish line. E: Apoptosis of erythrocyte in F3 homozygous *Tg*(*zgata1-g6pd*^*M118-144*^-*egfp*) transgenic zebrafish line.

## Notes

### Competing Interest Statement

The authors have declared no competing interest.

